# Effective Machine-Learning Assembly For Next-Generation Sequencing With Very Low Coverage

**DOI:** 10.1101/393116

**Authors:** Louis Ranjard, Thomas K. F. Wong, Allen G. Rodrigo

## Abstract

In short-read DNA sequencing experiments, the read coverage is a key parameter to successfully assemble the reads and reconstruct the sequence of the input DNA. When coverage is very low, the original sequence reconstruction from the reads can be difficult because of the occurrence of uncovered gaps. Reference guided assembly can then improve these assemblies. However, when the available reference is phylogenetically distant from the sequencing reads, the mapping rate of the reads can be extremely low. Some recent improvements in read mapping approaches aim at modifying the reference according to the reads dynamically. Such approaches can significantly improve the alignment rate of the reads onto distant references but the processing of insertions and deletions remains challenging. Here, we introduce a dynamic programming algorithm to update the reference sequence according to previously aligned reads. Substitutions, insertions and deletions are performed in the reference sequence dynamically. We evaluate this approach to assemble a western-grey kangaroo mitochondrial amplicon. Our results show that more reads can be aligned and that this method produces assemblies of length comparable to the truth while limiting error rate when classic approaches fail to recover the correct length. Our method allows us to assemble the first full mitochondrial genome for the western-grey kangaroo. Finally, we discuss how the core algorithm of this method could be improved and combined with other approaches to analyse larger genomic sequences.

## INTRODUCTION

De novo assembly algorithms classically use graph, de Bruijn or overlap-layout-consensus, to join short sequencing reads into longer contigs. However, when the short-reads coverage is very low, only short contigs can be reconstructed because of the occurrence of uncovered gaps in the sequence (Miller et al., 2010). In this case, the availability of a reference sequence can be beneficial to connect and order these contigs, an approach known as reference-guided assembly or homology-guided assembly (Rausch et al., 2009; Lischer and Shimizu, 2017). The reads are mapped onto this reference and a contig is constructed by taking the consensus of the short-reads at each position. However, some gaps in the mapping of the reads onto the reference may remain if the available reference is too distant phylogenetically from the sequence the short-reads originate from. This is because the short-reads that cannot, or can only partially, be mapped to the distant reference are discarded or trimmed. The information contained in the discarded or trimmed sequences of the reads is therefore lost. Hence, improvements in the alignments of the reads to the reference that are able to take advantage of this unexploited information should improve the assemblies.

Iterative referencing proposes to align all the reads to the reference and then update the reference sequence by calling the consensus of the reads. Once the reference has been updated, several additional iterations of read mapping/reference update can be performed to progressively improve the results (Otto et al., 2010; Tsai et al., 2010; Dutilh et al., 2011; Ghanayim, 2013; Hahn et al., 2013). Significant improvements in the mapping accuracy of the reads is achieved thanks to this approach (Břrinda et al., 2016). Subsequently, it has been shown that dynamic approaches can offer comparable improvements while performing less data processing, i.e. only requiring a single iteration of read mapping (Břrinda et al., 2016). In dynamic mapping, the reference is updated continuously as the reads are aligned onto it in an online fashion. Hence, the information obtained from the alignments of previous reads is used to map future reads. Dynamic strategies can be especially useful when the read sequences are highly divergent from the reference (Břrinda et al., 2016). However, the treatment of insertions and deletions (indels) remains a problem to dynamic mappers as the coordinates of the reads have to be continuously recalculated (Břrinda et al., 2016) with a new indexing of the reference.

Here, we assess how an online read aligner can improve the alignment of the reads when the reference is distant phylogenetically from the reads. This is a difficult task because, in this case, a large portion of the reads cannot be mapped to the reference. Using a machine learning approach, we present an algorithm that is able to dynamically perform substitutions and indels in the reference. The probability of each base at each position is learned from the past read alignments. A dynamic time warping algorithm uses these probability vectors directly to measure the edit distance between a read and the reference at the best alignment position. This is contrasting from previously proposed dynamic mapping approaches that record a counter for the different possible variants between the sequential updates of the reference (Břrinda et al., 2016). In the present method, the reference is updated after every read alignments. Note that our algorithm allows the reference to be updated with insertions and deletions at any position in the reference. We show that, because the reference sequence is continuously updated according to the alignment of the previous reads, the alignment of the read gradually improves. We demonstrate that this feature allows us to take advantage of distantly related reference sequence and improve the resulting short-reads assembly.

## RESULTS

In order to assess our method, we asked whether the improved read alignment provided by a dynamic approach results in better guided assemblies. We compared the assembly obtained from the dynamic aligner to classic assembly techniques. Briefly, we tested three assembly pipelines referred to as: **mapping**, mapping of the read to the reference followed by update of the reference; **learning**, dynamic time warping alignment of the reads with simultaneous machine learning approach to update the reference (Nucleoveq (Ranjard, 2018), see online Methods for details); **de novo**, reference-free assembly of the reads using a de Bruijn graph approach. Additionally, two hybrids approach were evaluated, the **de novo + mapping** and the **de novo + learning** pipelines where the contigs obtained by the de novo assembly of the reads are respectively mapped and aligned before updating the reference. A set of computer simulations was performed to compare the reconstructed sequence obtained by these strategies when coverage is very low (1 5 x) and with varying phylogenetic distances between the original sequence and the sequence used as reference.

We used sequencing short-reads obtained from a study of mitochondrial amplicons of the western-grey kangaroo, *Macropus fuliginosus* (Ranjard et al., 2018; Wong et al., 2018). Focusing on a 5, 000 bp amplicon allowed us to conduct extensive re-sampling of the reads. Published mitochondrial reference sequences from the following species were used as references: the eastern-grey kangaroo (*Macropus giganteus*, Genbank accession NC 027424), the swamp wallaby (*Wallabia bicolor*, Genbank accession KJ868164), the Tasmanian devil (*Sarcophilus harrisii*, Genbank accession JX475466) and the house mouse (*Mus musculus*, Genbank accession NC 005089). The computer simulations were performed using the most divergent amplicon (Amplicon 3) identified by Ranjard et al. (2018) which is located from position 11,756 to 16,897 in the eastern-grey kangaroo mitochondrial genome, total length of 5,130bp. This region contains the mitochondrial D-loop and, at the time of this study, the nucleotide sequence is not covered in the western-grey kangaroo mitochondrial genome (Genbank accession KJ868120). These species were chosen at increasing phylogenetic distance from the western-grey kangaroo (Table 1) but with no changes in their gene order. The homologous regions were selected in each species by aligning the amplicon sequence to each mitochondrial genome in Geneious version 10.2.4 (Kearse et al., 2012). Then, a region spanning from position 11,000 bp to 1,200 bp was used for each circular reference genome except the eastern-grey kangaroo. For the eastern-grey sequence the homologous amplicon region was used (Ranjard et al., 2018). This was done to reduced computational time while still keeping some part of the sequences located outside of the target region, i.e. from which the short-reads originate. The quality of the different assemblies was evaluated by using two statistics: first, the number of errors while aligning the reconstructed amplicon and the true western-grey kangaroo amplicon sequences; second, the length of the reconstructed sequence.

**Table 1.** The four different reference sequences used to guide the reconstruction of the western-grey kangaroo mitochondrial amplicon from short sequencing reads. The percentage identity is calculated on the homologous regions only, i.e. the non-aligned sections at the beginning and the end of the alignment are not taken into account.

| Species | Genbank accession | Length | Start position | End position | Percentage identity to western-grey amplicon |
| --- | --- | --- | --- | --- | --- |
| Eastern-grey kangaroo | NC_027424 | 5,186 | 11,756 | 16,897 | 91.4% |
| Swamp wallaby | KJ868164 | 7,075 | 11,000 | 1,200 | 86.8% |
| Tasmanian devil | JX475466 | 7,336 | 11,000 | 1,200 | 65.7% |
| House mouse | NC_005089 | 6,500 | 11,000 | 1,200 | 59.0% |

### Reference positions covered

The total read coverage in the reference was recorded for both the **mapping** and **learning** approaches to assess whether dynamic reference updates increases the reads alignment rate. As expected, the number of bases covered increases with the number of reads sampled (Figure 1). However, with distant reference sequences, i.e. the Tasmanian devil and the house mouse, the mapping rate of the reads is very low while the alignment rate is less affected by the increasing phylogenetic distance of the reference. Moreover, with these two species used as reference, the mapping rate remains low even though the depth of coverage increases. Generally, it appears that the variance in the mapping rate is higher than for the alignment rate.

**Figure 1.**
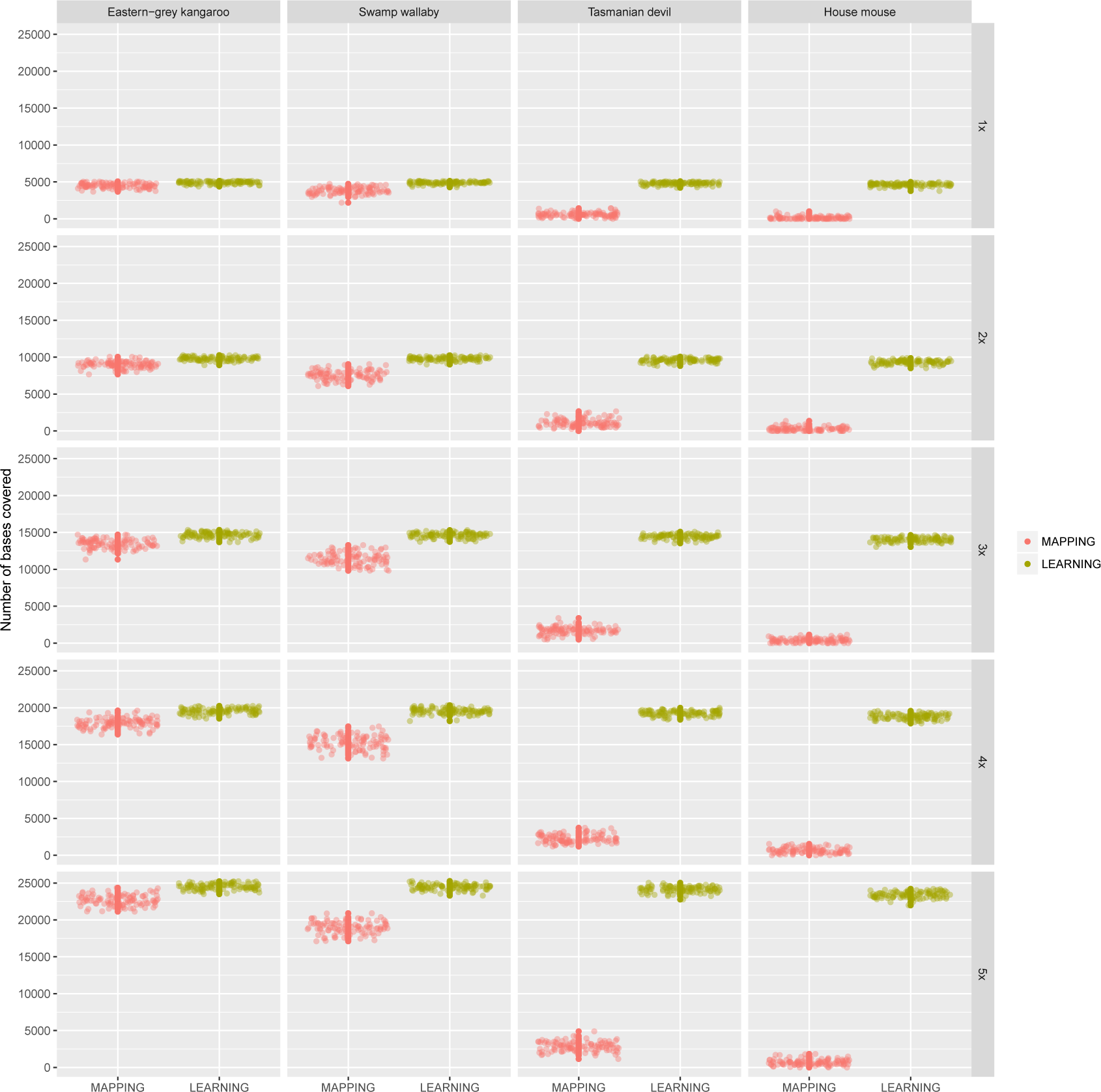
Realised coverage obtained by mapping (MAPPING) or aligning (LEARNING) sequencing reads to increasingly distant homologous reference sequences. The short-reads originate from a western-grey kangaroo amplicon.

### Assembly evaluation

A total of 2, 000 computer simulations were conducted. For coverage values ranging from 1× to 5×, the number of reads required to achieve such coverage was calculated and a corresponding subset of reads was randomly chosen among the full set. Then, for each of the four species reference sequence, the five pipelines were tested. A total of 100 replicates was performed for each setting. To compute the number of errors and length of the reconstructed sequence statistics, the pairwise alignment was computed using the Needleman-Wunsch algorithm with affine gap penalty scheme, the NUC44 scoring matrix and null gap penalties at the end of the sequences. The non-aligned sequences at the beginning and at the end of the alignment were discarded and the remaining sequence length was reported for comparisons between pipelines. The number of errors was computed as the Hamming distance between the remaining aligned sequences.

Overall, the **learning** approaches offered the best compromise between limiting the error rate and recovering the true length of the amplicon sequence (Figure 2). In all simulation settings, the de Bruijn graph assemblies (**de novo assembly**) achieved a very low error rate. On the other hand, this approach was only able to generate relatively short assemblies compared to the other pipelines (Figure 2). However, with increasing coverage the length of the de novo assembled contigs increased confirming the suitability of de Bruijn graph based methods for assembling short-reads when the depth of coverage is high. When using distant references (Tasmanian devil and the house mouse), the hybrid approaches (**de novo + mapping** and **de novo + learning**) produced less errors than the same algorithms used on the raw reads. However, when using more closely related sequences as references, the **de novo + mapping** method produced more errors than the **mapping** pipeline. This is putatively the consequence of the low coverage of the de novo assembly of the reads, i.e. the **de novo** only generated very short contigs. On the other hand, the **de novo + learning** and **learning** generated similar amount of errors with closely related reference sequences used as guides. With more distant reference sequences, the **de novo + learning** produced less errors than the **learning** pipeline. While both pipelines benefit from an increase in read coverage, the **de novo + learning** returned the lowest amount of errors with distant references.

**Figure 2.**
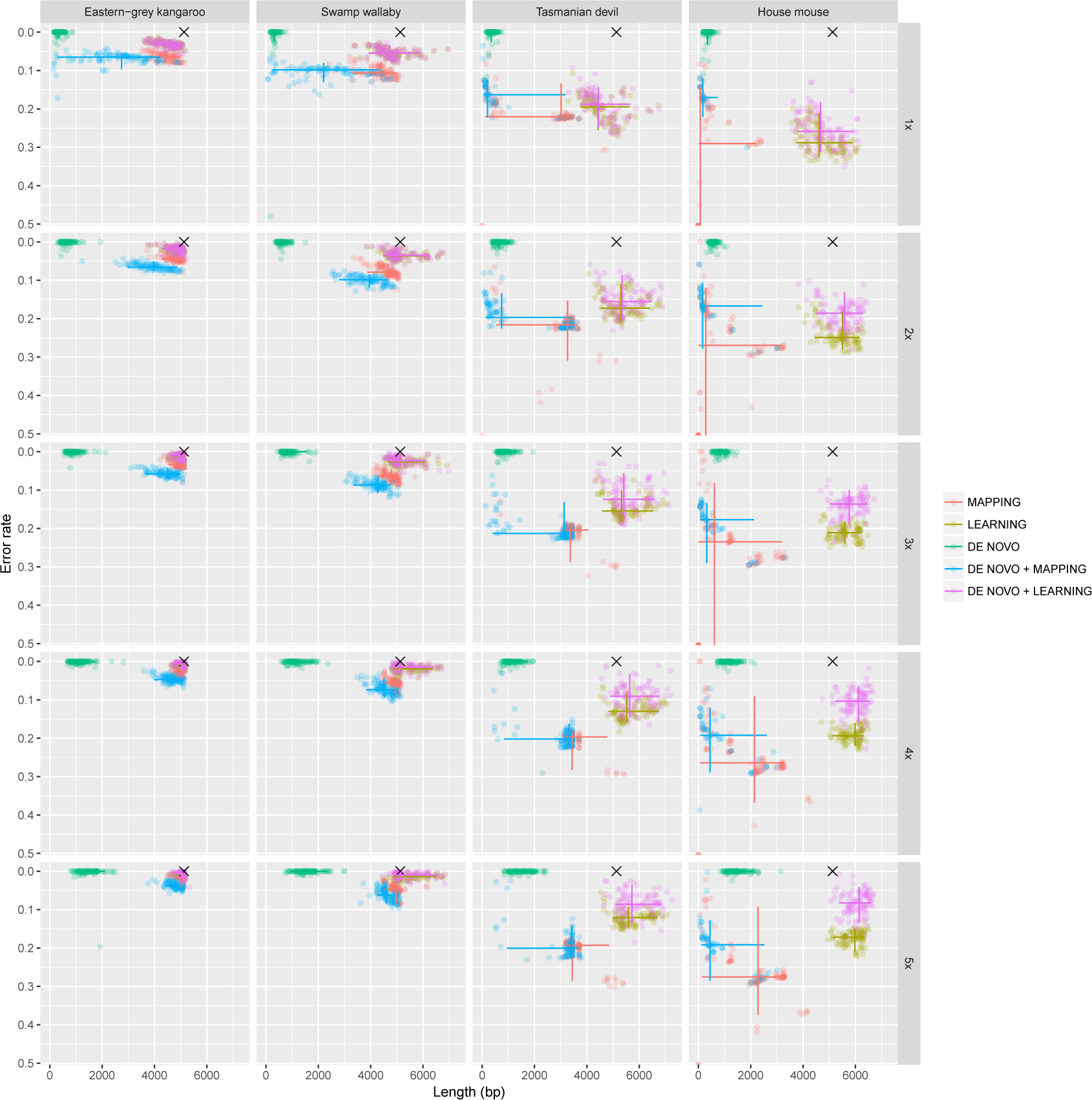
Number of errors and length in nucleotide of the reconstructed amplicon for each bioinformatic pipeline and simulation settings. The 95% intervals are shown as solid lines for each method along both dimensions (reconstructed amplicon length and error rate).

When the reference sequence was chosen phylogenetically close to the reads sequence, i.e. eastern-grey kangaroo and swamp wallaby, and the coverage was set to 5×, all pipelines, except **de novo assembly**, generated assemblies of comparable length from the truth. With decreasing coverage, the reconstructed sequence length also decreased for all methods. This is particularly noticeable for approaches that use mapping of the reads as the mapping rate strongly decreases with increasing phylogenetic distance of the reference (Figure 1). On the other hand, the two methods that use dynamic programming to align the reads were able to reconstruct sequences of length comparable to the western-grey amplicon using distant reference (Figure 2). It is noticeable that in these cases the variance of both the length and the error rate for the mapping-based pipelines is comparatively very high. This is highly likely to be the consequence of the higher variance in the mapping rate for these pipelines and it may indicate that the mapping-based methods are more sensitive to a non-uniform coverage of the re-sampled reads.

### Comparison to iterative referencing

Additionally, an iterative mapping approach was implemented by repeating the **mapping** pipeline five times using the updated reference obtained at the previous iteration. This approach was tested with the Tasmanian devil reference sequence at coverage 5× as it is expected that the best improvements would be obtained with higher coverage. As expected iterative mapping improved the sequence reconstruction (Table 2). Each additional iteration of the mapping of the reads allowed the error rate to decrease as more reads could be mapped. However, the improvements were limited. After five iterations, the error rate and the length of the reconstructed sequence were still worse than the ones obtained with the **de novo + learning** pipeline (Table 2). Similar limited improvements were obtained using the other reference sequences and coverage values (data not shown).

**Table 2.**
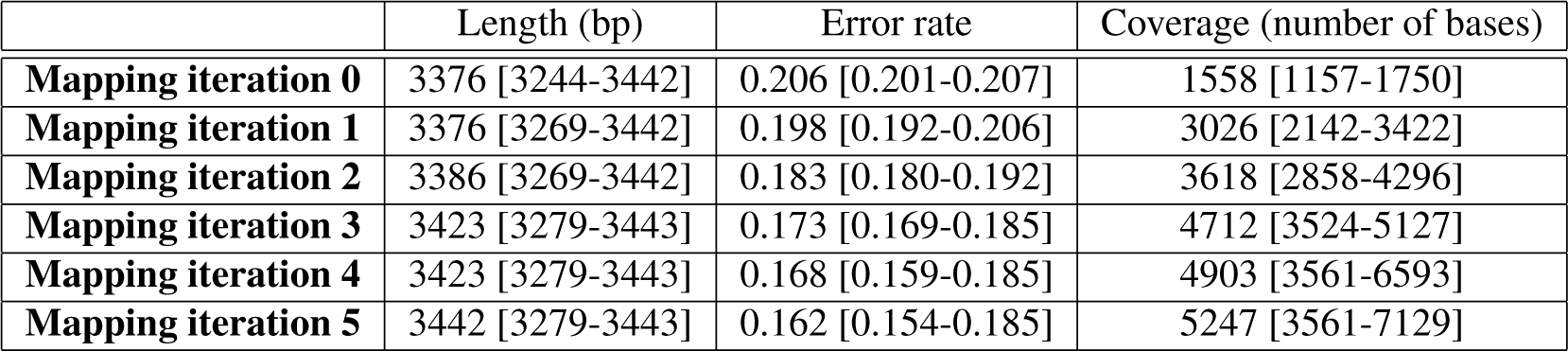
Iterative mapping lowers the error rate and the length of the reconstructed sequences. The median and the first and third quartiles are indicated for each statistic, the coverage was 5 with the Tasmanian devil used as reference sequence.

### Assembly of *Macropus fuliginosus* mitochondrial genome

To demonstrate the applicability of the method, a full mitochondrial genome was assembled from short-reads using a sister species reference sequence. At the time of this study, the western-grey kangaroo mitochondrial genome is only partial and lacks the hyper variable region (Genbank accession KJ868120) (Ranjard et al., 2018). We used our method to reconstruction the full mitochondrial genome of the individual identified as “KA” in Ranjard et al. (2018). First, the partial mitochondrial genome of the western-grey kangaroo was completed using the eastern-grey kangaroo reference (Genbank accession NC 027424) generating an hybrid full genome template. The sequencing reads generated from three western-grey kangaroo mitochondrial amplicons, of length 4641bp, 4152bp and 5140bp (83% of the genome, Ranjard et al. (2018)), were then aligned to this reference template using Nucleoveq. One of the amplicon fully spans the missing region in the western-grey kangaroo mitochondrial genome reference. Reads were sub-sampled so that to obtain a coverage of 5× . Because the coverage was low, ten iterations were conducted to insure that the reference was fully covered by randomly sampled reads.

The ten replicates of the mitochondrial genome assembly with an average of 99% identity. Visual inspections of the alignment of the replicates showed that these differences occurred in regions with no coverage. The consensus sequence of the ten replicates was compared to the high coverage assembly of the mitochondrial assembly from Ranjard et al. (2018). As expected, some errors were observed at the beginning or end of the three mitochondrial amplicons. Because the short-read coverage was extremely low in these regions, it was very unlikely that the sub sampling of the reads retrieved these sequences. A new mitochondrial genome was generated by correcting the consensus sequence with the high coverage information. The newly assembled western-grey mitochondrial genome was annotated in Geneious version 10.2.4 (Kearse et al., 2012) using the eastern-grey kangaroo mitochondrial genome as a reference. The western-grey complete mitochondrial genome is on Genbank under accession number MH717106.

## DISCUSSION

By iteratively aligning short sequencing reads and updating the reference sequence, we were able to improve the reconstruction of the read sequence, resulting in assemblies of comparable length to the truth while limiting the number of errors. The improvement of this dynamic alignment method over the graph- or the mapping-based approaches tested here can be explained by two factors. First, the alignment rate is higher when using dynamic programming over the Burrows-Wheeler transform approach used for mapping the reads. Second, the progressive modifications of the reference, as reads are aligned onto it, facilitate the alignment of the following reads because the reference is continuously pulled closer to the reads sequence (Břrinda et al., 2016). This is particularly useful when only a phylogenetically distant reference sequence is available for a reference-guided assembly. Actually, our results showed that the static mapping of the reads is not possible when the reference is too distant from the reads, as demonstrated by a very low mapping rate.

The drawback of our dynamic programming method for read alignment is memory usage. The memory required to build the alignment matrix *M* precludes the direct usage of this method for large genome assemblies. While our approach is relevant to small genome assemblies, e.g. mitochondrial, supplementary work would be required to adapt this approach to large genome read alignments. For example, while it is not possible to directly align the reads to a large genome, a first search could help identify short windows, i.e. few thousands bases, in the reference sequence where the reads could then be aligned more accurately by our algorithm. In the current implementation of the method, it is optionally possible to take advantage of the known mapping positions of the reads by passing a mapping file as argument. This technique can massively reduce the memory requirements as only a window of specified size around these positions will be considered for performing the alignment. Our algorithm could also be combined with other methods to find the potential locations of each read in the genome prior to performing the alignments. The seed-based algorithm used by Blast (Altschul et al., 1990) or some kmer-based seed searches (Liao et al., 2013; Břrinda et al., 2015) are obvious candidates. However, when the reference sequence is distant from the reads, it is not possible to initially map all the reads onto it. It is therefore inevitable to re-align or re-map these reads once the reference has been partially updated.

Our method improves previous dynamic reference building approaches in that it allows the reference to be updated with insertions and deletions. Previously, Liao et al. (2013) proposed a seed and vote approach to locate indels. Břrinda et al. (2016) proposed a dynamic mapping approach where the reference is iteratively updated with the read sequences but indels were not fully supported (Břrinda et al., 2017). Our method not only locates but also aligns and corrects the reference sequence with indels, facilitating further the subsequent read alignments. This approach comes at the computational cost of realigning each read onto the reconstructed reference. However, in our algorithm each read is treated independently and the updates of the reference are only performed according to the information from one read at a time. This is different from graph-based and iterative referencing methods that need all reads to be aligned before calling the variants. As a consequence, parallelization may be used to distribute batch of reads to be analysed independently prior to merging the several assemblies.

The threshold limit for performing insertions and deletions was set to be equal to the learning rate (see online Methods). Therefore, indels will not be performed when the read alignment is poor. However, there is no particular reasons to use this value and other values could be used based on other statistics. Preliminary tests (data not shown) indicated that this value nevertheless returned best assemblies. Similarly, the indels costs was set to equal the maximum possible distance between a pair of nucleotide vectors. Preliminary tests using grid search showed that similar results were obtained while varying their values (data not shown). However, this hyper-parameters could also be set to depend on some other parameters measured on the data and further investigations could be conducted to explore these possibilities.

Finally, the learning rate hyper-parameter was set to depend on the alignment distance. Classically in machine learning algorithms, the learning rate is set to decay through the learning process (Ranjard et al., 2015; Ruder, 2016). Conversely, in our algorithm, it is expected that the rate will increase as the reference sequence gets closer to the reads. Alternative learning rate schedules could be tested, for example cyclic methods as proposed by Smith (2015) for training deep neural networks. Moreover, we only considered one epoch for learning, i.e. one iteration over the full set of reads. In other words, the total read set is only seen once to learn the amplicon sequence. Because the reads are chosen in a random order, the assembled sequence will potentially be different between distinct runs of the algorithm and there is no guarantee to converge on the best assembly. Performing the learning over multiple epochs could potentially improve the convergence among runs at the cost of processing time.

## ONLINE METHODS

### Learning from dynamic programming alignment of the reads to the reference

In essence, the algorithm consists in aligning the reads to the reference using dynamic time warping. Then, an “average” sequence of the aligned region is computed from the best path of the local free-ends alignment (Ranjard and Ross, 2008). This approach was originally designed to perform unsupervised clustering of bioacoustic sequences (Ranjard et al., 2017). In this work, a similar algorithm is implemented to analyse nucleotide sequences: each nucleotide position in a sequence is represented as a four elements vector, the Voss representation (Voss, 1992), encoding the probability of each base according to previously aligned reads. This numerical representation of DNA sequence is appropriate for the comparison of DNA sequences (Mendizabal-Ruiz et al., 2017) and their classification (Mendizabal-Ruiz et al., 2018). In molecular biology, similar algorithm has been applied to the clustering of amino acid sequences (Olshen et al., 2005) where vector quantization is used to estimate the probability density of amino acids. In the area of genomic signal processing, dynamic time warping approaches have been successful at classifying various representations of genomic data (Legrand et al., 2008; Skutkova et al., 2013, 2015; Loose et al., 2016).

We consider two sequences of nucleotide vectors, a reference *F* = *f*_1_ *… f_l_* and a read *R* = *r* _1_ *… r_n_*, respectively representing the reference sequence of length *l* and a read of length *n* aligned onto it. The vectors *f_x_*, where 1 ≤ *x ≤ l*, and *r_y_*, where 1 ≤ *y ≤ n*, represent the probability vectors of each nucleotide at position *x* in the reference and position *y* in the read, respectively. Through a statistical learning process and vector quantization, the reference sequence vectors are updated according to the sequencing read nucleotides. Ultimately, the goal is to reconstruct, i.e. assemble, the original sequence *S* from which the reads come from.

A probability vector *r_y_* is calculated according to the quality scores of each base at position *y* and, at initialisation, all *f_x_* are only made of binary vectors defined by the reference sequence. Additionally, a “persistence” vector *P* = *p* _1_*...p_l_*, where *p_i_* for 1 ≤ *i ≤ l* are initialised all to 1, is updated when indels occur for each nucleotide position in the reference. The distance between a pair of nucleotide vectors is defined as

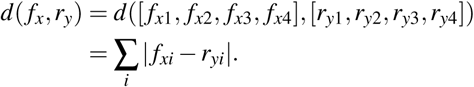

A dynamic programming approach is used to align the reads to the reference sequence. Let *M* (*x, y*) the minimum edit distance over all possible suffixes of the reference from position 1 to *x* and the read from position 1 to *y*.

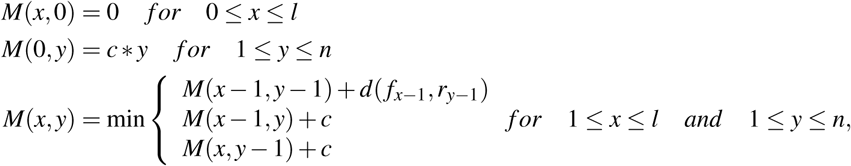

with the insertion/deletion cost is *c* = 1. The three elements correspond to three edit operations: insertion, deletion and substitution. The value in *e_FR_* = *min*_1*≤x≤l*_ *M*(*x, n*) therefore consists in an edit distance between the read and the reference vector sequences of nucleotide vectors. It is then normalised by the length of the read to obtain a read “edit rate”, *ê*_*FR*_.

The optimal path is traced back and, at each position, the new reference vector is updated. In case of a substitution, *f_x_* = *w f_x_* + (1 — *w*)*r_y_* with a learning rate *w* (see below). In cases of deletions or insertions, the *f_x_* remains unchanged but the corresponding position in the persistence vector decreases or increases by an amount equal to (1 — *w*), respectively. Then, the persistence value is assessed against a threshold: if *p_x_ >* 1 + *w* or *p_x_ <* 1 — *w*, then an insertion or a deletion is performed at the position *x* in the reference sequence. For insertions, the inserted nucleotide vector is initialised to the same value *r_y_* which is the nucleotide probability vector on the position *y* of the read *r* aligned to the inserted position in the reference. All the reads are chosen in random order and sequentially aligned to the reference sequence according to this procedure (Figure 3).

**Figure 3.**
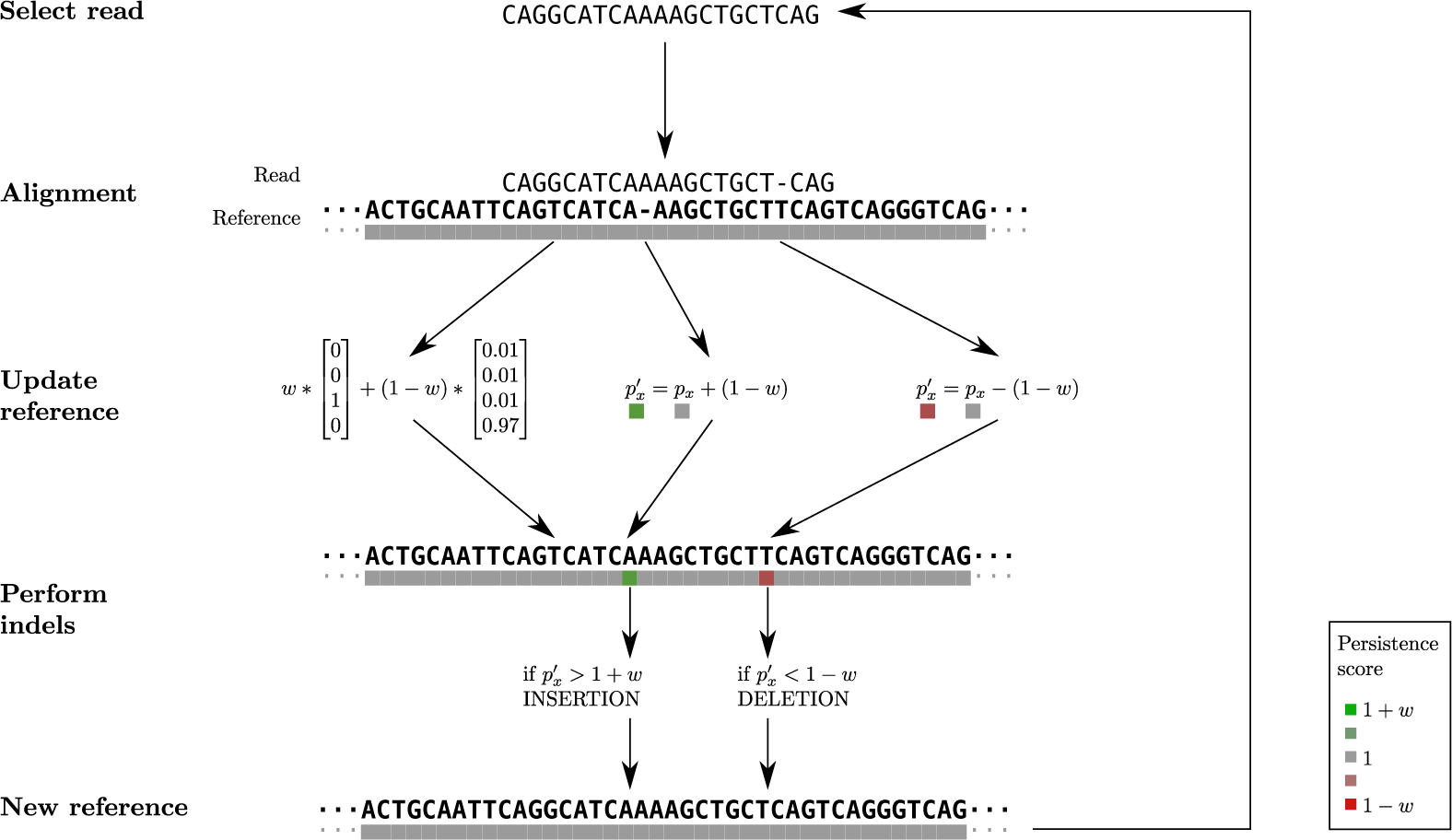
Overview of the algorithm. Reads are taken in random order and iteratively aligned to the reference. After each alignment, the reference sequence is updated according to the learning rate *w*, which is proportional to the normalised edit distance between the read and the reference. In this case, there is one substitution between the reference of the read; the read has a *G* with Phred quality score of 15 while the reference is *T* . One deletion and one insertion are treated thanks to a persistence vector. The persistence value *p_•_* indicates the tendency of a base to be inserted or deleted at each position in the reference. This value can trigger indels update in the reference when it goes beyond a threshold.

### Learning rate

The learning rate (1 — *w*) is set to depend on the edit rate and governs how much the reference is updated. For low values of (1 − *w*) the reference mostly remains unmodified. When the distance between the read and the reference is low, there is high certainty in the positioning of the read onto the reference. Therefore, the learning rate can be increased to facilitate the update of the reference toward the sequence of the read. On the other hand, when the alignment of the read is more difficult, i.e. high edit distance, the learning rate is set to a low value so that the reference is only slightly updated and misalignments or errors in the read sequence are not affecting the learning process.

Computer simulations were conducted in order to determine the distribution of the edit distances between reads and increasingly divergent reference sequences. First, a nucleotide sequence of length *u*(500,5000) was generated by randomly choosing nucleotides with 50% GC content. A read sequence of length 150 was generated by randomly choosing a position in the original sequence and using an error rate of 1% with the errors uniformly distributed along the sequence. Then, mutations were introduced in the original sequence, at a rate of 1, 5, 10, 30, 50 %, and single nucleotide indels were introduced at a rate of 10%. Additionally, random reference sequences of similar length were generated to build a random distribution of the distance. The process was repeated 1, 000 times (Figure 4).

**Figure 4.**
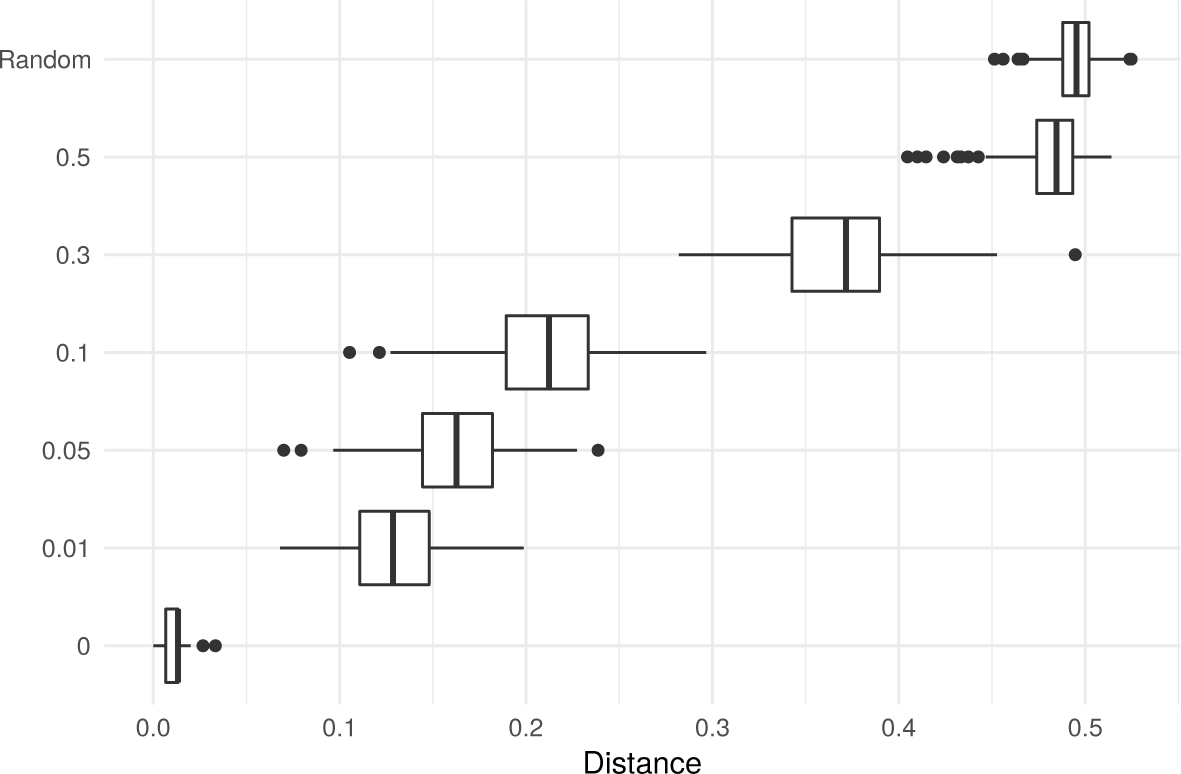
Distribution of the normalised edit distance between reads and increasingly distant reference sequences. The mutation rate of the reference sequence is indicated on the y-axis. The top row (Random) shows the distribution of the edit distance when reads were aligned to randomly generated nucleotide sequences. For the lowest row, the reads were aligned to their original sequence and the departure from 0 of the edit distance only results from the simulated sequencing errors.

From the empirical distributions of the distance (Figure 4), the learning rate was determined to be equal to 0.95 when the distance is below 0.05, which corresponds to the range of distances expected due to sequencing errors. It is set to 0.05 when the distance is above 0.35, i.e. the distance expected when the read and the reference sequence have less than 70% sequence similarity. Between normalised edit distances of 0.05 and 0.95, the rate was set to linearly increase, i.e. 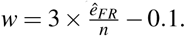.

### Five assembly pipelines

First, the whole set of reads, average coverage of ~2, 000, was mapped to the eastern-grey kangaroo to determine the western-grey kangaroo mitochondrial sequence for the amplicon (see Ranjard et al. (2018) for details). Then, five different bioinformatic pipelines were tested at lower coverage. At first, the reads were preprocessed before running each pipeline: Illumina adapters and low quality bases were removed (Trimmomatic version 0.36, Bolger et al. (2014)) using a sliding window of 15 nucleotides, with steps of four bases and the resulting reads below length 36 were discarded. Additionally, kmer error correction was performed using Tadpole (BBMap version 37.95, Brian Bushnell). The five assembly pipelines (Figure 5) are described below:

1. **Mapping** was performed using Bowtie2 version 2.2.6 (Langmead and Salzberg, 2012). Both “local” alignment with “soft trimmed” and “end-to-end” alignment of the reads were tested. In general, local alignment resulted in higher alignment rates and was therefore used in all simulations. Once the reads were aligned to the reference, Samtools version 1.5 (Li, 2011) was used to order the reads. Freebayes version 1.1.0 (Garrison and Marth, 2012) then allowed us to identify variants. Calls with high probability to be false positive, Phred score *<* 20, were removed with Vcffilter (Vcflib version 1.0.0) (Garrison, 2016). The consensus sequence was generated using Bcftools version 1.6 (Li, 2011) by applying the alternative variants to the reference sequence. Finally, the uncovered parts at the beginning and at the end of the reference were removed.
2. **Learning** consisted in iteratively aligning the reads and dynamically updating the reference according to the machine learning approach previously described, the algorithm is implemented in Nucleoveq (Ranjard, 2018). For these simulations, all the reads were aligned to the reference and no prior information about the mapping position was utilised to perform read alignments. At the end of the learning process, the uncovered regions located at the beginning and end of the reference were truncated to generate the final assembly.
3. **De novo assembly** was done with Trinity version 2.4.0 (Grabherr et al., 2011), using a kmer size of 17 and setting the minimum contig length to 100 so that assembly could be performed when coverage was very low. After assembly, the longest contig was selected for evaluation.
4. **De novo + Mapping** consisted in mapping all the **de novo assembly** contigs obtained from Trinity to the reference in an effort to connect them into a longer sequence. The same approach as for **mapping** pipeline was used to generate the consensus.
5. **De novo + Learning** consisted in feeding all the **de novo assembly** contigs obtained from Trinity to our machine learning algorithm. The same steps as for the above **learning** pipeline were performed while regarding the contigs instead of the reads as input.

**Figure 5.**
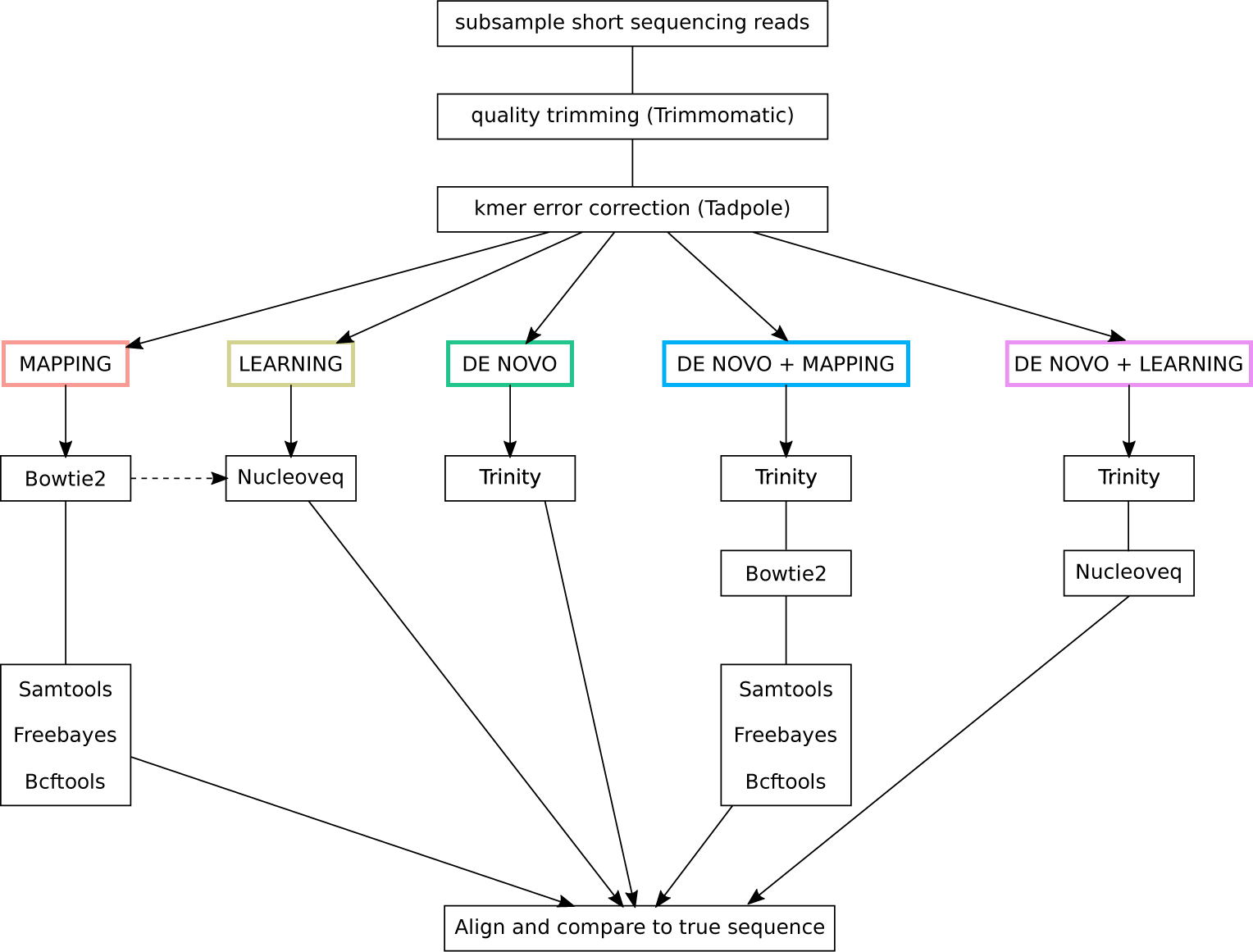
Five bioinformatic pipelines for assembly. Dashed-line: it is possible to pass a priori mapping position of the reads to Nucleoveq to decrease memory requirements and speed up computation (option not used in the reported comparisons).

